# Interpretability of Multivariate Brain Maps in Brain Decoding: Definition and Quantification

**DOI:** 10.1101/047522

**Authors:** Seyed Mostafa Kia

## Abstract

Brain decoding is a popular multivariate approach for hypothesis testing in neuroimaging. Linear classifiers are widely employed in the brain decoding paradigm to discriminate among experimental conditions. Then, the derived linear weights are visualized in the form of multivariate brain maps to further study the spatio-temporal patterns of underlying neural activities. It is well known that the brain maps derived from weights of linear classifiers are hard to interpret because of high correlations between predictors, low signal to noise ratios, and the high dimensionality of neuroimaging data. Therefore, improving the interpretability of brain decoding approaches is of primary interest in many neuroimaging studies. Despite extensive studies of this type, at present, there is no formal definition for interpretability of multivariate brain maps. As a consequence, there is no quantitative measure for evaluating the interpretability of different brain decoding methods. In this paper, first, we present a theoretical definition of interpretability in brain decoding; we show that the interpretability of multivariate brain maps can be decomposed into their reproducibility and representativeness. Second, as an application of the proposed definition, we formalize a heuristic method for approximating the interpretability of multivariate brain maps in a binary magnetoencephalography (MEG) decoding scenario. Third, we pro pose to combine the approximated interpretability and the performance of the brain decoding into a new multi-objective criterion for model selection. Our results for the MEG data show that optimizing the hyper-parameters of the regularized linear classifier based on the proposed criterion results in more informative multivariate brain maps. More importantly, the presented definition provides the theoretical background for quantitative evaluation of interpretability, and hence, facilitates the development of more effective brain decoding algorithms in the future.

## 1 Introduction

Understanding the mechanisms of the brain has been a crucial topic throughout the history of science. Ancient Greek philosophers envisaged different functionalities for the brain ranging from cooling the body to acting as the seat of the rational soul and the center of sensation [1]. Modern cognitive science, emerging in the 20th century, provides better insight into the brain’s functionality. In cognitive science, researchers usually analyze recorded brain activity and behavioral parameters to discover the answers of *where, when*, and *how* a brain region participates in a particular cognitive process.

To answer the key questions in cognitive science, scientists often employ mass-univariate hypothesis testing methods to test scientific hypotheses on a large set of independent variables [2, 3]. Mass-univariate hypothesis testing is based on performing multiple tests, e.g., t-tests, one for each unit of the neuroimaging data, i.e., independent variables. The high spatial and temporal granularity of the univariate tests provides fair level of interpretability. On the down side, the high dimensionality of neuroimaging data requires a large number of tests that reduces the sensitivity of these methods after multiple comparison correction. Although some techniques such as the non-parametric cluster-based permutation test [4, 5] offer more sensitivity because of the cluster assumption, they still experience low sensitivity to brain activities that are narrowly distributed in time and space [2, 6]. The multivariate counterparts of mass-univariate analysis, known generally as multivariate pattern analysis (MVPA), have the potential to overcome these deficits. Multivariate approaches are capable of identifying complex spatio-temporal interactions between different brain areas with higher sensitivity and specificity than univariate analysis [7], especially in group analysis of neuroimaging data [8].

*Brain decoding* [9] is an MVPA technique that delivers a model to predict the mental state of a human subject based on the recorded brain signal. There are two potential applications for brain decoding: 1) brain-computer interfaces (BCIs) [10, 11], and 2) multivariate hypothesis testing [12]. In the first case, a brain decoder with maximum prediction power is desired. In the second case, in addition to the prediction power, extra information on the spatio-temporal nature of a cognitive process is desired. In this study, we are interested in the second application of brain decoding that can be considered a multivariate alternative for mass-univariate hypothesis testing.

In brain decoding, generally, linear classifiers are used to assess the relation between independent variables, i.e., features, and dependent variables, i.e., cognitive tasks [13, 14, 15]. This assessment is performed by solving a linear optimization problem that assigns weights to each independent variable. Currently, brain decoding is the gold standard in multivariate analysis for functional magnetic resonance imaging (fMRI) [16, 17, 18, 19] and magnetoencephalogram/electroencephalogram (MEEG) studies [20, 21, 22, 23, 24, 25, 26]. It has been shown that brain decoding can be used in combination with brain encoding [27] to infer the causal relationship between stimuli and responses [28].

*Brain mapping* [29] is a higher form of neuroimaging that assigns pre-computed quantities, e.g., univariate statistics or weights of a linear classifier, to the spatio-temporal representation of neuroimaging data. In MVPA, brain mapping uses the learned parameters from brain decoding to produce brain maps, in which the engagement of different brain areas in a cognitive task is visualized. Intuitively, the interpretability of a brain decoder refers to the level of information that can be reliably derived by an expert from the resulting maps. From the neuroscientific perspective, a brain map is considered interpretable if it enables the scientist to answer *where, when*, and *how* questions.

Typically, a trained classifier is a black box that predicts the label of an unseen data point with some accuracy. Valverde-Albacete and Peláez-Moreno [30] experimentally showed that in a classification task optimizing only classification error rate is insufficient to capture the transfer of crucial information from the input to the output of a classifier. It is also shown by Ramdas et al. [31] that in the case of data with small sample size using the classification accuracy as a test statistic for two sample testing should be performed with extra cautious. Beside these limitations of classification accuracy in inference, and considering the fact that the best predictive model might not be the most informative one [32]; a classifier, taken alone, only answers the question of *what* is the most likely label of a given unseen sample [33]. This fact is generally known as knowledge extraction gap [34] in the classification context. Thus far, many efforts have been devoted to filling the knowledge extraction gap of linear and non-linear data modeling methods in different areas such as computer vision [35], signal processing [36], chemometrics [37], bioinformatics [38], and neuroinformatics [39].

Despite the theoretical advantages of MVPA, its practical application to inferences regarding neuroimaging data is limited primarily by a lack of interpretability [40, 41, 42]. Therefore, improving the interpretability of linear brain decoding and associated brain maps is a primary goal in the brain imaging literature [43]. The lack of interpretability of multivariate brain maps is a direct consequence of low signal-to-noise ratios (SNRs), high dimensionality of whole-scalp recordings, high correlations among different dimensions of data, and cross-subject variability [15, 44, 45, 14, 46, 47, 48, 49, 50, 51, 52, 41]. At present, two main approaches are proposed to enhance the interpretability of multivariate brain maps: 1) introducing new metrics into the model selection procedure and 2) introducing new penalty terms for regularization to enhance stability selection.

The first approach to improving the interpretability of brain decoding concentrates on the model selection procedure. Model selection is a procedure in which the best values for the hyper-parameters of a model are determined [14]. The selection process is generally performed by considering the generalization performance, i.e., the accuracy, of a model as the decisive criterion. Rasmussen et al. [53] showed that there is a trade-off between the spatial reproducibility and the prediction accuracy of a classifier; therefore, the reliability of maps cannot be assessed merely by focusing on their prediction accuracy. To utilize this finding, they incorporated the spatial reproducibility of brain maps in the model selection procedure. An analogous approach, using a different definition of spatial reproducibility, is proposed by Conroy et al. [54]. Beside spatial reproducibility, the stability of the classifiers [55] is another criterion that is used in combination with generalization performance to enhance the interpretability. For example, [56, 57] showed that incorporating the stability of models into cross-validation improves the interpretability of the estimated parameters (by linear models).

The second approach to improving the interpretability of brain decoding focuses on the underlying mechanism of regularization. The main idea behind this approach is two-fold: 1) customizing the regularization terms to address the ill-posed nature of brain decoding problems (where the number of samples is much less than the number of features) [58, 50] and 2) combining the structural and functional prior knowledge with the decoding process so as to enhance stability selection. Group Lasso [59] and total-variation penalty [60] are two effective methods using this technique [61, 62]. Sparse penalized discriminant analysis [63], group-wise regularization [7], randomized Lasso [47], smoothed-sparse logistic regression [64], total-variation L1 penalization [65, 66], the graph-constrained elastic-net [67, 68], and randomized structural sparsity [69] are examples of brain decoding methods in which regularization techniques are employed to improve stability selection, and thus, the interpretability of brain decoding.

Recently, taking a new approach to the problem, Haufe et al. questioned the interpretability of weights of linear classifiers because of the contribution of noise in the decoding process [70, 39, 71]. To address this problem, they proposed a procedure to convert the linear brain decoding models into their equivalent generative models. Their experiments on the simulated and fMRI/EEG data illustrate that, whereas the direct interpretation of classifier weights may cause severe misunderstanding regarding the actual underlying effect, their proposed transformation effectively provides interpretable maps. Despite the theoretical soundness, the major challenge of estimating the empirical covariance matrix of the small sample size neuroimaging data [72] limits the practical application of this method.

In spite of the aforementioned efforts to improve the interpretability of brain decoding, there is still no formal definition for the interpretability of brain decoding in the literature. Therefore, the interpretability of different brain decoding methods are evaluated either qualitatively or indirectly (i.e., by means of an intermediate property). In qualitative evaluation, to show the superiority of one decoding method over the other (or a univariate map), the corresponding brain maps are compared visually in terms of smoothness, sparseness, and coherency using already known facts (see, for example, [47, 73]). In the second approach, important factors in interpretability such as spatio-temporal reproducibility are evaluated to indirectly assess the interpretability of results (see, for example, [46, 53, 54, 74]). Despite partial effectiveness, there is no general consensus regarding the quantification of these intermediate criteria. For example, in the case of spatial reproducibility, different methods such as correlation [53, 74], dice score [46], or parameter variability [39, 54] are used for quantifying the stability of brain maps, each of which considers different aspects of local or global reproducibility.

With the aim of filling this gap, our contribution is three-fold: 1) Assuming that the true solution of brain decoding is available, we present a theoretical definition of the interpretability. Furthermore, we show that the interpretability can be decomposed into the reproducibility and the representativeness of brain maps. 2) As a proof of the concept, we propose a practical heuristic based on event-related fields for quantifying the interpretability of brain maps in MEG decoding scenarios. 3) Finally, we propose the combination of the interpretability and the performance of the brain decoding as a new Pareto optimal multi-objective criterion for model selection. We experimentally show that incorporating the interpretability into the model selection procedure provides more reproducible, more neurophysiologically plausible, and (as a result) more interpretable maps.

## 2 Methods

### 2.1 Notation and Background

Let 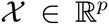 be a manifold in Euclidean space that represents the input space and 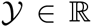 be the output space, where 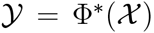. Then, let 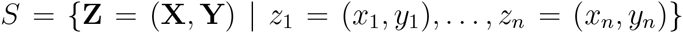 be a training set of *n* independently and identically distributed (iid) samples drawn from the joint distribution of 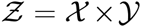 based on an unknown Borel probability measure *ρ*. In the neuroimaging context, X indicates the trials of brain recording, e.g., fMRI, MEG, or EEG signals, and Y represents the experimental conditions or dependent variables. The goal of brain decoding is to find the function 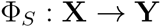 as an estimation of the ideal function 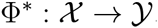.

As is a common assumption in the neuroimaging context, we assume the true solution of a brain decoding problem is among the family of linear functions 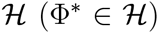. Therefore, the aim of brain decoding reduces to finding an empirical approximation of 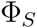, indicated by 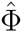, among all 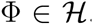. This approximation can be obtained by estimating the predictive conditional density 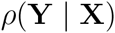 by training a parametric model 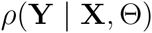 (i.e., a likelihood function), where Θ denotes the parameters of the model. Alternatively, Θ can be estimated by solving a risk minimization problem:

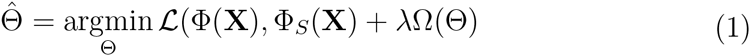

where 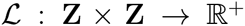 is the loss function, 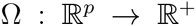 is the regularization term, and λ is a hyper-parameter that controls the amount of regularization. There are various choices for Ω, each of which reduces the hypothesis space 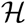 to 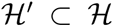 by enforcing different prior functional or structural constraints on the parameters of the linear decoding model (see, for example, [75, 76, 60, 77]). The amount of regularization A is generally decided using cross-validation or other data perturbation methods in the model selection procedure.

In the neuroimaging context, the estimated parameters of a linear decoding model 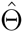 can be used in the form of a brain map so as to visualize the discriminative neurophysiological effect. Although the magnitude of 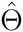 is affected by the dynamic range of data and the level of regularization, it has no effect on the predictive power and the interpretability of maps. On the other hand, the direction of 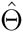 affects the predictive power and contains information regarding the importance of and relations among predictors. This type of relational information is very useful when interpreting brain maps in which the relation between different spatio-temporal independent variables can be used to describe how different brain regions interact over time for a certain cognitive process. Therefore, we refer to the normalized parameter vector of a linear brain decoder in the unit hyper-sphere as a multivariate brain map (MBM); we denote it by 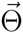 where 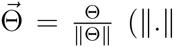 represents the 2-norm of a vector).

As shown in Eq. 1, learning occurs using the sampled data. In other words, in the learning paradigm, we attempt to minimize the loss function with respect to 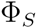 (and not 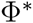) [78]. Therefore, all of the implicit assumptions (such as linearity) regarding 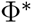 might not hold on 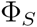, and vice versa (see the supplementary material for a simple illustrative example). The *irreducible error* ɛ is the direct consequence of sampling; it sets a lower bound on the error, where we have:

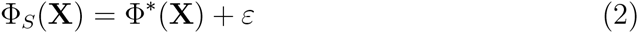

The distribution of ɛ dictates the type of loss function 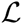 in Eq. 1. For example, assuming a Gaussian distribution with mean 0 and variance 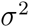 for ɛ implies the least squares loss function [79].

### 2.2 Interpretability of Multivariate Brain Maps: Theoretical Definition

In this section, we present a theoretical definition for the interpretability of linear brain decoding models and their associated MBMs. Our definition of interpretability is based on two main assumptions: 1) the brain decoding problem is linearly separable; 2) its *unique* and neurophysiologically *plausible*^1^ solution, i.e., 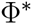, is available.

Consider a linearly separable brain decoding problem in an ideal scenario where *ɛ* = 0 and *rank*(*X*) = *p*. In this case, 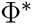 is linear and its parameters 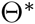 are unique and plausible. The unique parameter vector 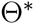 can be computed as follows:

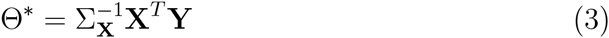

Using 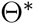 as the reference, we define the *strong-interpretability* of an MBM as follows:

**Definition 1**. An MBM 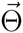 associated with a linear function Φ is “strongly-interpretable” if and only if 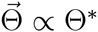.

It can be shown that, in practice, the estimated solution of a linear brain problem (using Eq. 1) is not strongly-interpretable because of the inherent limitations of neuroimaging data, such as uncertainty [80] in the input and output space (*ɛ* ≠ 0), the high dimensionality of data (*n* << *p*), and the high correlation between predictors (rank(X) < p). With these limitations in mind, even though in practice the solution of linear brain decoding is not strongly-interpretable, one can argue that some are more interpretable than others. For example, in the case in which 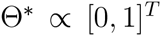, a linear classifier where 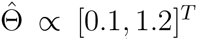 can be considered more interpretable than a linear classifier where 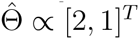. This issue raises the following question:

**Problem 1**. Let *S_1_*,…, *S_m_* be *m* perturbed training sets drawn from *S* via a certain perturbation scheme such as jackknife, bootstrapping [81], or cross-validation [82]. Assume 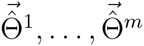 are *m* MBMs of a certain Φ (estimated using Eq. 1 for certain 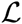, Ω, and λ) on the corresponding perturbed training sets. How can we quantify the proximity of Φ to the strongly-intrepretable solution of brain decoding problem 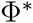?

To answer this question, considering the uniqueness and the plausibility of 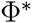 as the two main characteristics that convey its strong-interpretability, we define the interpretability as follows:

**Definition 2**. Let 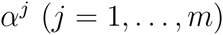 be the angle between 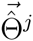 and 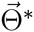. The “interpretability” 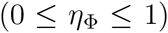 of the MBM derived from a linear function Φ is defined as follows:

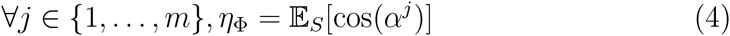

Empirically, the interpretability is the mean of cosine similarities between 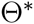 and MBMs derived from different samplings of the training set. In addition to the fact that employing cosine similarity is a common method for measuring the similarity between vectors, we have another strong motivation for this choice. It can be shown that, for large values of p, the distribution of the dot product in the unit hyper-sphere, i.e., the cosine similarity, converges to a normal distribution with 0 mean and variance of 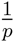, i.e., 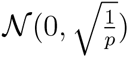. Due to the small variance for a large enough p values, any similarity value that is significantly larger than zero represents a meaningful similarity between two high dimensional vectors (see the supplementary material for more details about the distribution of cosine similarity).

In what follows, we demonstrate how the definition of interpretability is geometrically related to the uniqueness and plausibility characteristics of the true solution to brain decoding problem.

### 2.3 Interpretability Decomposition into Reproducibility and Representativeness

An alternative approach toward quantifying the interpretability is to assess separately its uniqueness and neurophysiological plausibility. In this section, we firstly define the reproducibility and representativeness as measures for quantifying the uniqueness and neurophysiological plausibility of brain decoding model, respectively. Then we show how these definitions are related to the definition of interpretability.

The high dimensionality and the high correlations between variables are two inherent characteristics of neuroimaging data that negatively affect the uniqueness of the solution of a brain decoding problem. Therefore, a certain configuration of hyper-parameters may result different estimated parameters on different portions of data. Here, we are interested in assessing this variability as a measure for uniqueness. Let 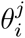 be the ith (*i* = 1,…,*p*) element of an MBM estimated on the *j*th (*j* = 1,…,*m*) perturbed training set. We first define the *main multivariate brain map* as follows:

**Definition 3**. The “main multivariate brain map” 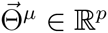 of a linear function Φ is defined as the sum of estimated MBMs 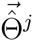 (*j* = 1,…,*m*) on the perturbed training sets *S^j^* in the unit hyper-sphere:

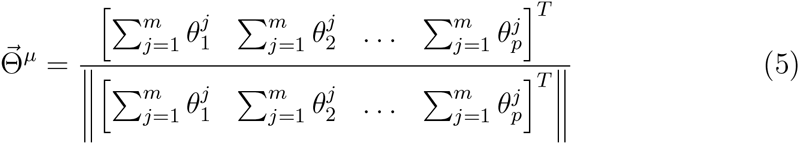

The definition of 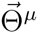 is analogous to the main prediction of a learning algorithm [83]; it provides a reference for quantifying the reproducibility of an MBM:

**Definition 4**. Let 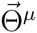 be the main multivariate brain map of Φ. Then, let α*^j^* be the angle between 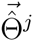 and 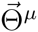. The “reproducibility” 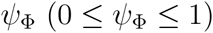 of an MBM derived from a linear function Φ is defined as follows:

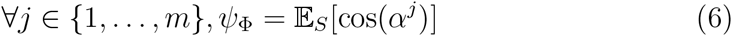

In fact, reproducibility provides a measure for quantifying the dispersion of MBMs, computed over different perturbed training sets, from the main multivariate brain map.

On the other hand, the coherency between the main multivariate brain map of a decoder and the true solution can be employed as a measure for the plausibility of a model. We refer to this coherency as the *representativeness* of an MBM:

**Definition 5**. Let 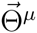 be the main multivariate brain map of Φ. The “representativeness” 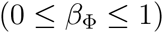 is defined as the cosine similarity between 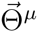 and 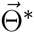:

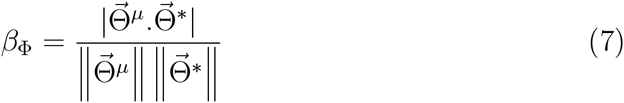

The following proposition shows the relationship between the presented definitions for reproducibility, representativeness, and the interpretability:

**Proposition 1**. 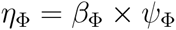.

See Appendix D for a proof. Proposition 1 indicates the interpretability can be decomposed into the representativeness and the reproducibility of a decoding model.

### 2.4 A Heuristic for Practical Quantification of Interpretability in Time-Domain MEG decoding

In practice, it is impossible to evaluate the interpretability, as 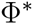 is unknown. In this study, to provide a practical proof of the mentioned theoretical concepts, we propose the use of contrast event-related fields (cERFs) of MEG data as neurophysiological plausible heuristics for 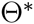 in a binary MEG decoding scenario in the time domain.

The EEG/MEG data are a mixture of several simultaneous stimulus-related and stimulus-unrelated brain activities. In general, unrelated-stimulus brain activities are considered as Gaussian noise with zero mean and variance 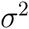. One popular approach to canceling the noise component is to compute the average of multiple trials. It is expected that the average will converge to the true value of the signal with a variance of 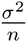. The result of the averaging process is generally known as ERF in the MEG context; separate interpretation of different ERF components can be performed [84]^1^.

Assume 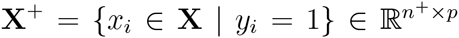 and 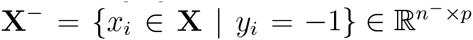. Then, the cERF brain map 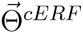 is computed as follows:

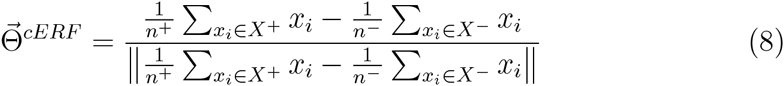

Using the core theory presented in [39], it can be shown that cERF is the equivalent generative model for the least squares solution in a binary time-domain MEG decoding scenario (see Appendix A). Using 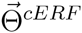 as a heuristic for 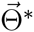, the representativeness can be approximated as follows:

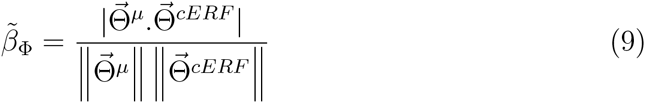

Where 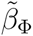 is an approximation of 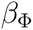 and we have:

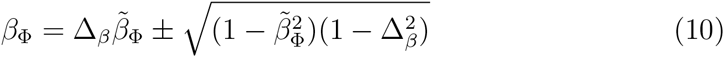

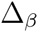 represents the cosine similarity between 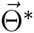 and 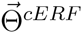 (see Figures B.8 and Appendix B). If 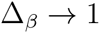 then 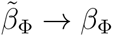.

In a similar manner, 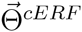 can be used to heuristically approximate the interpretability as follows:

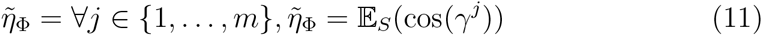

where 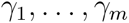 are the angles between 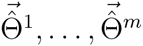 and 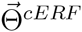. The following equality represents the relation between *η* and 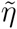 (see Figures C.9 and Appendix C).

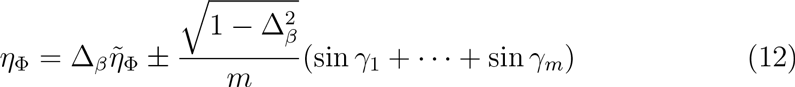

Again, if 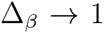 then 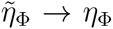. Notice that 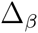 is independent of the decoding approach used; it only depends on the quality of the heuristic. It can be shown that 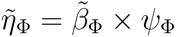.

Eq. 12 shows that the choice of heuristic has a direct effect on the approximation of interpretability and that an inappropriate selection of the heuristic yields a very poor estimation of interpretability because of the destructive contribution of 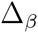. Therefore, the choice of heuristic should be carefully justified based on accepted and well-defined facts regarding the nature of the collected data (see the supplementary material for the experimental investigation of the limitations of the proposed heuristic).

### 2.5 Incorporating the Interpretability into Model Selection

The procedure for evaluating the performance of a model so as to choose the best values for hyper-parameters is known as *model selection* [85]. This procedure generally involves numerical optimization of the model selection criterion. The most common model selection criterion is based on an estimator of generalization performance, i.e., the predictive power. In the context of brain decoding, especially when the interpretability of brain maps matters, employing the predictive power as the only decisive criterion in model selection is problematic in terms of interpretability [86, 53, 54]. Here, we propose a multi-objective criterion for model selection that takes into account both prediction accuracy and MBM interpretability.

Let 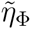 and 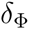 be the approximated interpretability and the generalization performance of a linear function Φ, respectively. We propose the use of the *scalarization* technique [87] for combining 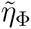 and 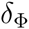 into one scalar 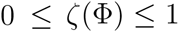 as follows:

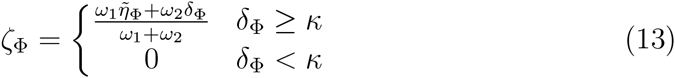

where 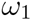 and 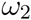 are weights that specify the level of importance of the interpretability and the performance, respectively. *κ* is a threshold on the performance that filters out solutions with poor performance. In classification scenarios, *κ* can be set by adding a small safe interval to the chance level of classification.

It can be shown that the hyper-parameters of a model Φ are optimized based on 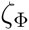 are Pareto optimal [88]. In other words, there exist no other 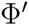 for which we obtain both 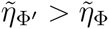 and 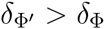. We expect that optimizing the hyper-parameters based on 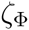, rather only 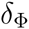, yields more informative MBMs.

### 2.6 Experimental Materials

#### 2.6.1 Toy Dataset

To illustrate the importance of integrating the interpretability of brain decoding with the model selection procedure, we use simple 2-dimensional toy data presented in [39]. Assume that the true underlying generative function 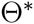 is defined by

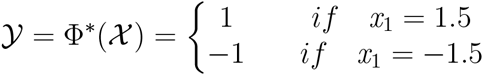

where 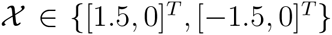; and *x*_1_ and *x*_2_ represent the first and the second dimension of the data, respectively. Furthermore, assume the data is contaminated by Gaussian noise with co-variance 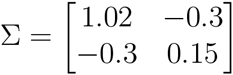. Figure 1 shows the distribution of the noisy data.

**Figure 1.**
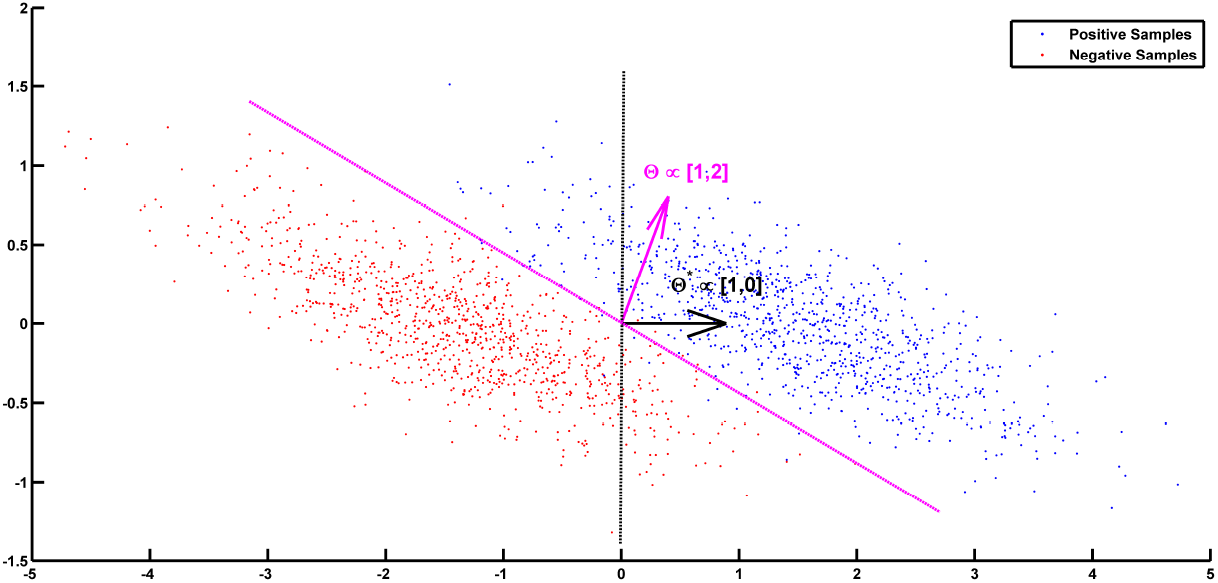
Noisy samples of toy data. The black line shows the true separator based on the generative model 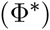. The magenta line shows the most accurate classification solution. Because of the contribution of noise, any interpretation of the parameters of the most accurate classifier yields a misleading conclusion with respect to the true underlying phenomenon [39].

#### 2.6.2 MEG Data

We use the MEG dataset presented in [89]^1^. The dataset was also used for the DecMeg2014 competition^2^. In this dataset, visual stimuli consisting of famous faces, unfamiliar faces, and scrambled faces are presented to 16 subjects and fMRI, EEG, and MEG signals are recorded. Here, we are only interested in MEG recordings. The MEG data were recorded using a VectorView system (Elekta Neuromag, Helsinki, Finland) with a magnetometer and two orthogonal planar gradiometers located at 102 positions in a hemispherical array in a light Elekta-Neuromag magnetically shielded room.

Three major reasons motivated the choice of this dataset: 1) It is publicly available. 2) The spatio-temporal dynamic of the MEG signal for face vs. scramble stimuli has been well studied. The event-related potential analysis of EEG/MEG shows that N170 occurs 130 - 200*ms* after stimulus presentation and reflects the neural processing of faces [90, 89]. Therefore, the N170 component can be considered the ground truth for our analysis. 3) In the literature, non-parametric mass-univariate analysis such as cluster-based permutation tests is unable to identify narrowly distributed effects in space and time (e.g., an N170 component) [2, 6]. These facts motivate us to employ multivariate approaches that are more sensitive to these effects.

As in [51], we created a balanced face vs. scrambled MEG dataset by randomly drawing from the trials of unscrambled (famous or unfamiliar) faces and scrambled faces in equal number. The samples in the face and scrambled face categories are labeled as 1 and -1, respectively. The raw data is high-pass filtered at 1*Hz*, down-sampled to 250*Hz*, and trimmed from 200ms before the stimulus onset to 800ms after the stimulus. Thus, each trial has 250 time-points for each of the 306 MEG sensors (102 magnetometers and 204 planar gradiometers)^1^. To create the feature vector of each sample, we pooled all of the temporal data of 306 MEG sensors into one vector (i.e., we have *p* = 250 × 306 = 76500 features for each sample). Before training the classifier, all of the features are standardized to have a mean of 0 and standard-deviation of 1.

### 2.7 Classification and Evaluation

In all experiments, a least squares classifier with L1-penalization, i.e., Lasso [75], is used for decoding. Lasso is a very popular classification method in the context of brain decoding, mainly because of its sparsity assumption. The choice of Lasso helps us to better illustrate the importance of including the interpretability in the model selection. Lasso solves the following optimization problem:

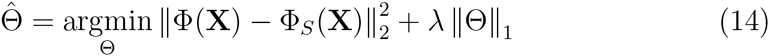

where λ is the hyper-parameter that specifies the level of regularization. Therefore, the aim of the model selection is to find the best value for λ. Here, we try to find the best regularization parameter value among λ = {0.001, 0.01, 0.1,1,10, 50,100, 250, 500,1000,5000,10000,15000, 25000, 50000}.

We use the out-of-bag (OOB) [91, 92] method for computing 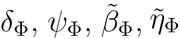, and 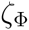 for different values of λ. In OOB, given a training set (X,Y), *m* replications of bootstrap [81] are used to create perturbed training sets (we set *m* = 50)^2^. In all of our experiments, we set 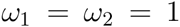 and *k* = 0.6 in the computation of 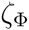. Furthermore, we set 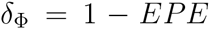 where EPE indicates the expected prediction error; it is computed using the procedure explained in Appendix E. Employing OOB provides the possibility of computing the bias and variance of the model as contributing factors in EPE.

To investigate the behavior of the proposed model selection criterion, we benchmark it against the commonly used performance criterion in the single-subject decoding scenario. Assuming (X*_i_*,Y*_i_*) for *i* = 1,…, 16 are MEG trial/label pairs for subject *i*, we separately train a Lasso model for each subject to estimate the parameter of the linear function 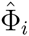, where 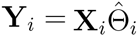. Let 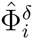 and 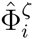 represent the optimized solution based on 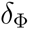 and 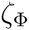, respectively. We denote the MBM associated with 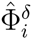 and 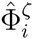 by 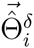 and 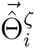, respectively. Therefore, for each subject, we compare the resulting decoders and MBMs computed based on these two model selection criteria.

## 3 Results

### 3.1 Performance-Interpretability Dilemma: A Toy Example

In the definition of 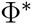 on the toy dataset discussed in Section 2.6.1, *x_1_* is the decisive variable and *x_2_* has no effect on the classification of the data into target classes. Therefore, excluding the effect of noise and based on the theory of the maximal margin classifier [93, 94], 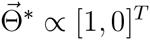 is the true solution to the decoding problem. By accounting for the effect of noise and solving the decoding problem in (X, Y) space, we have 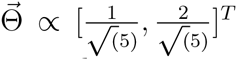 as the parameter of the linear classifier. Although the estimated parameters on the noisy data yield the best generalization performance for the noisy samples, any attempt to interpret this solution fails, as it yields the wrong conclusion with respect to the ground truth (it says *x*_2_ has twice the influence of *x*_1_ on the results, whereas it has no effect). This simple experiment shows that the most accurate model is not always the most interpretable one, primarily because the contribution of the noise in the decoding process [39]. On the other hand, the true solution of the problem 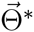 does not provide the best generalization performance for the noisy data.

To illustrate the effect of incorporating the interpretability in the model selection, a Lasso model with different λ values is used for classifying the toy data. In this case, because 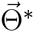 is known, the exact value of interpretability can be computed using Eq. 4. Table 1 compares the resultant performance and interpretability from Lasso. Lasso achieves its highest performance 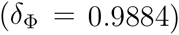 at λ = 10 with 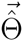 ∝ [0.4636,0.8660]*^T^* (indicated by the magenta line in Figure 1). Despite having the highest performance, this solution suffers from a lack of interpretability 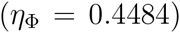. By increasing λ, the interpretability improves so that for λ = 500,1000 the classifier reaches its highest interpretability by compensating for 0.06 of its performance. Our observation highlights two main points:

1. In the case of noisy data, the interpretability of a decoding model is incoherent with its performance. Thus, optimizing the parameter of the model based on its performance does not necessarily improve its interpretability. This observation confirms the previous finding by Rasmussen et al. [53] regarding the trade-off between the spatial reproducibility (as a measure for the interpretability) and the prediction accuracy in brain decoding.
2. If the right criterion is used in the model selection, employing proper regularization technique (sparsity prior, in this case) leads to more interpretability for the decoding models.

**Table 1.**
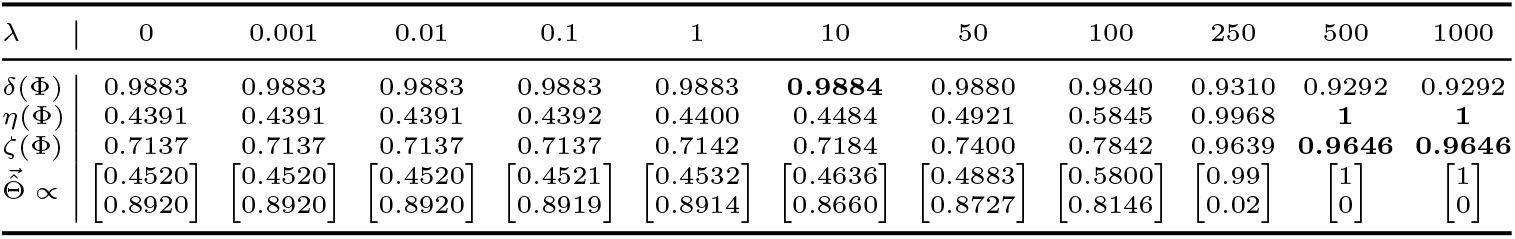
Comparison between 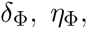 and 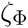 for different λ values on the toy 2D example shows the performance-interpretability dilemma, in which the most accurate classifier is not the most interpretable one.

### 3.2 Mass-Univariate Hypothesis Testing on MEG Data

Results show that non-parametric mass-univariate analysis is unable to detect narrowly distributed effects in space and time (e.g., an N170 component) [2, 6]. To illustrate the advantage of the proposed decoding framework for spotting these effects, we performed a non-parametric cluster-based permutation test [5] on our MEG dataset using Fieldtrip toolbox [95]. In a single subject analysis scenario, we considered the trials of MEG recordings as the unit of observation in a between-trials experiment. Independent-samples t-statistics are used as the statistics for evaluating the effect at the sample level and to construct spatio-temporal clusters. The maximum of the cluster-level summed t-value is used for the cluster level statistics; the significance probability is computed using a Monte Carlo method. The minimum number of neighboring channels for computing the clusters is set to 2. Considering 0.025 as the two-sided threshold for testing the significance level and repeating the procedure separately for magnetometers and combined-gradiometers, no significant result is found for any of the 16 subjects. This result motivates the search for more sensitive (and, at the same time, more interpretable) alternatives for hypothesis testing.

### 3.3 Single-Subject Decoding on MEG Data

In this experiment, we aim to compare the multivariate brain maps of brain decoding models when 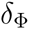 and 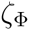 are used as the criteria for model selection. Figure 2(a) represents the mean and standard-deviation of the performance and interpretability of Lasso across 16 subjects for different A values. The performance and interpretability curves further illustrate the performance-interpretability dilemma in the single-subject decoding scenario in which increasing the performance delivers less interpretability. The average performance across subjects is improved when λ approaches 1, but on the other side, the reproducibility and the representativeness of models declines significantly [see Figure 2(b)].

**Figure 2.**
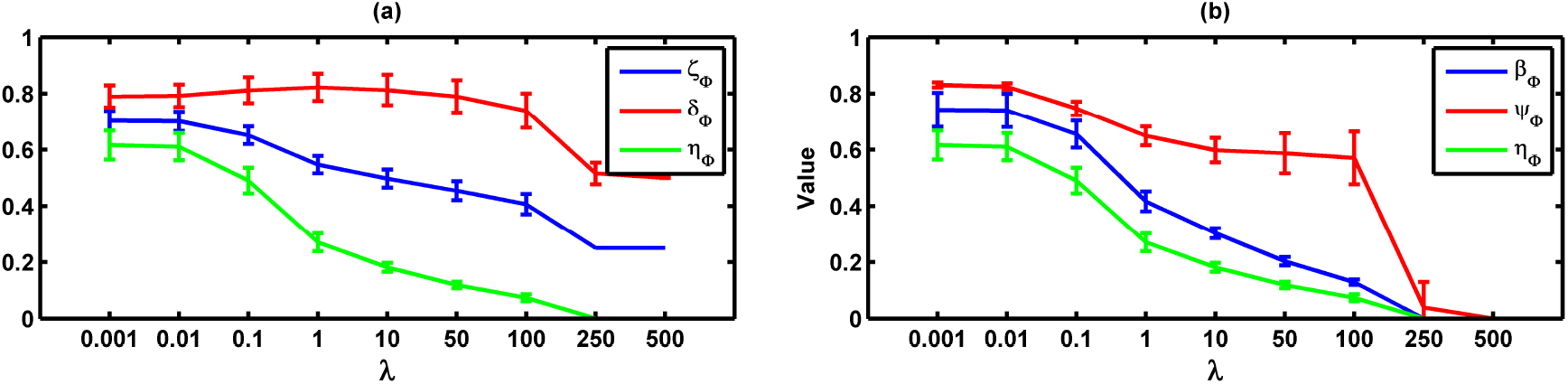
(a) Mean and standard-deviation of the performance, interpretability, and plausibility of Lasso over 16 subjects. The performance and interpretability become incoherent as λ increases. (b) Mean and standard-deviation of the reproducibility, representativeness, and interpretability of Lasso over 16 subjects. The interpretability declines because of the decrease in both reproducibility and representativeness.

One possible reason behind the performance-interpretability dilemma is illustrated in Figure 3. The figure shows the mean and standard deviation of bias, variance, and EPE of Lasso across 16 subjects. The plot proposes that the effect of variance is overwhelmed by bias in the computation of EPE, where the best performance (minimum EPE) at λ = 1 has the lowest bias, its variance is higher than for λ = 0.001,0.01,0.1. While this tiny increase in the variance is not reflected in EPE but Figure 2(b) shows a significant effect on the reproducibility.

**Figure 3.**
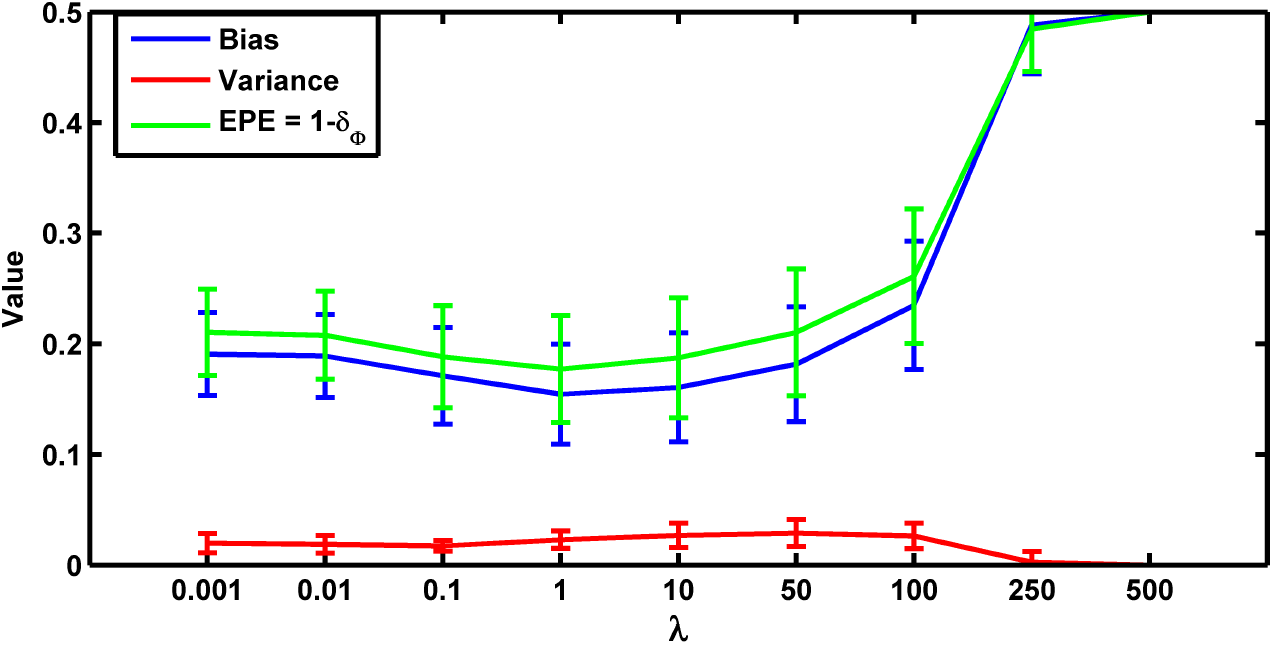
Mean and standard-deviation of the bias, variance, and EPE of Lasso over 16 subjects. The effect of variance on the EPE is overwhelmed by bias.

Table 2 summarizes the performance, reproducibility, representativeness, and interpretability of 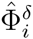 and 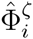 for 16 subjects. The average result over 16 subjects shows that employing 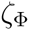 instead of 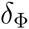 in model selection provides significantly higher reproducibility, representativeness, and (as a result) interpretability compensating for 0.04 of performance.

**Table 2.**
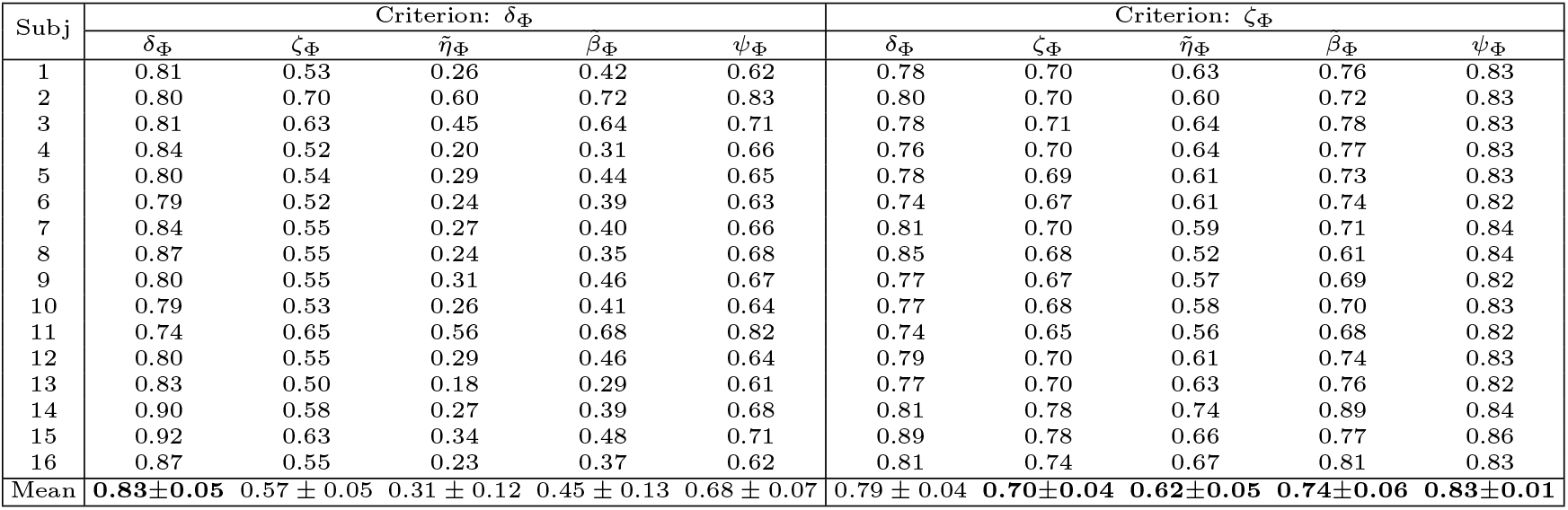
The performance, reproducibility, representativeness, and interpretability of 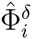 and 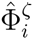 over 16 subjects.

These results are further analyzed in Figure 4 where 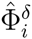 and 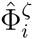 are compared subject-wise in terms of their performance and interpretability. The comparison shows that adopting 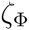 instead of 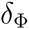 as the criterion for model selection yields significantly better interpretable models by compensating a negligible degree of performance in 14 out of 16 subjects. Figure 4(a) shows that employing 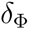 provides on average slightly higher accurate models (Wilcoxon rank sum test p-value= 0.012) across subjects (0.83 ± 0.05) than using 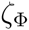 (0.79±0.04). On the other side, Figure 4(b) shows that employing 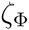 and compensating by 0.04 in the performance provides (on average) substantially higher (Wilcoxon rank sum test p-value= 5.6 × 10^−6^) interpretability across subjects (0.62 ± 0.05) compared to 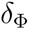 (0.31 ± 0.12). For example, in the case of subject 1 (see table 2), using 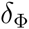 in model selection to select the best λ value for the Lasso yields a model with 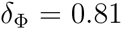 and 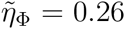. In contrast, using 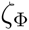 delivers a model with 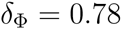 and 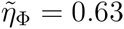.

**Figure 4.**
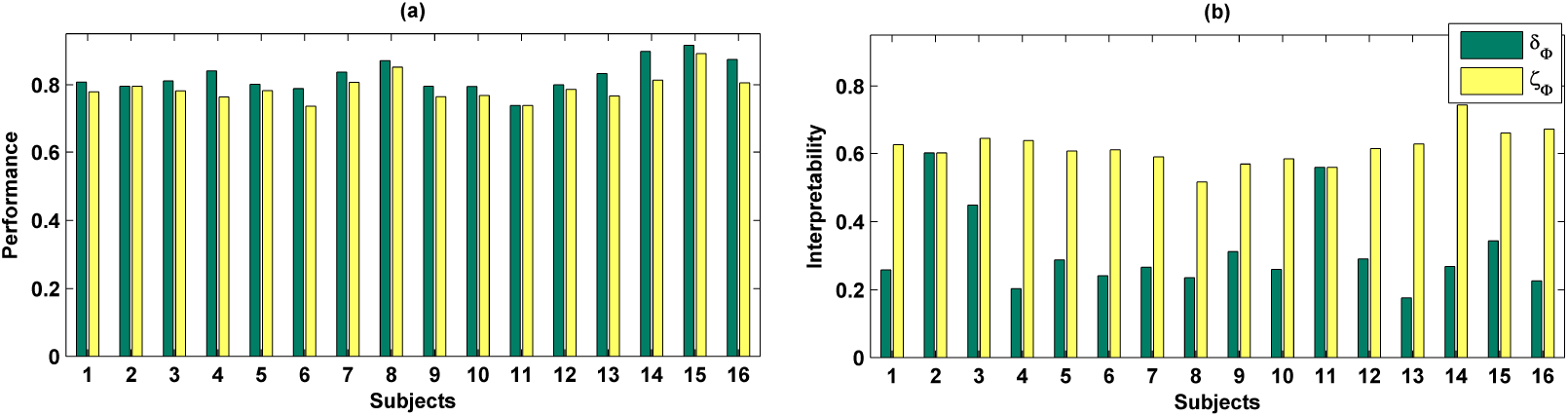
a) Comparison between performance of 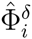 and 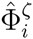. Adopting 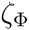 instead of 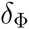 in model selection yields (on average) 0.04 less accurate classifiers over 16 subjects. b) Comparison between interpretability of 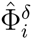 and 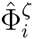. Adopting 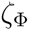 instead of 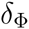 in model selection yields on average 0.31 more interpretable classifiers over 16 subjects.

The advantage of the exchange between the performance and the interpretability can be seen in the quality of MBMs. Figure 5a and 5b show 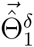 and 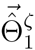 of subject 1, i.e., the spatio-temporal multivariate maps of the Lasso models with maximum values of 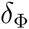 and 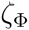, respectively. The maps are plotted for 102 magnetometer sensors. In each case, the time course of weights of classifiers associated with the MEG2041 and MEG1931 sensors are plotted. Furthermore, the topographic maps represent the spatial patterns of weights averaged between 184*ms* and 236*ms* after stimulus onset^1^. While 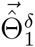 is sparse in time and space, it fails to accurately represent the spatio-temporal dynamic of the N170 component. Furthermore, the multi-collinearity problem arising from the correlation between the time course of the MEG2041 and MEG1931 sensors causes extra attenuation of the N170 effect in the MEG1931 sensor. Therefore, the model is unable to capture the spatial pattern of the dipole in the posterior area. In contrast, 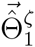 represents the dynamic of the N170 component in time (see Figure 6). In addition, it also shows the spatial pattern of two dipoles in the posterior and temporal areas. In summary, 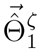 suggests a more representative pattern of the underlying neurophysiological effect than 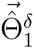.

**Figure 5.**
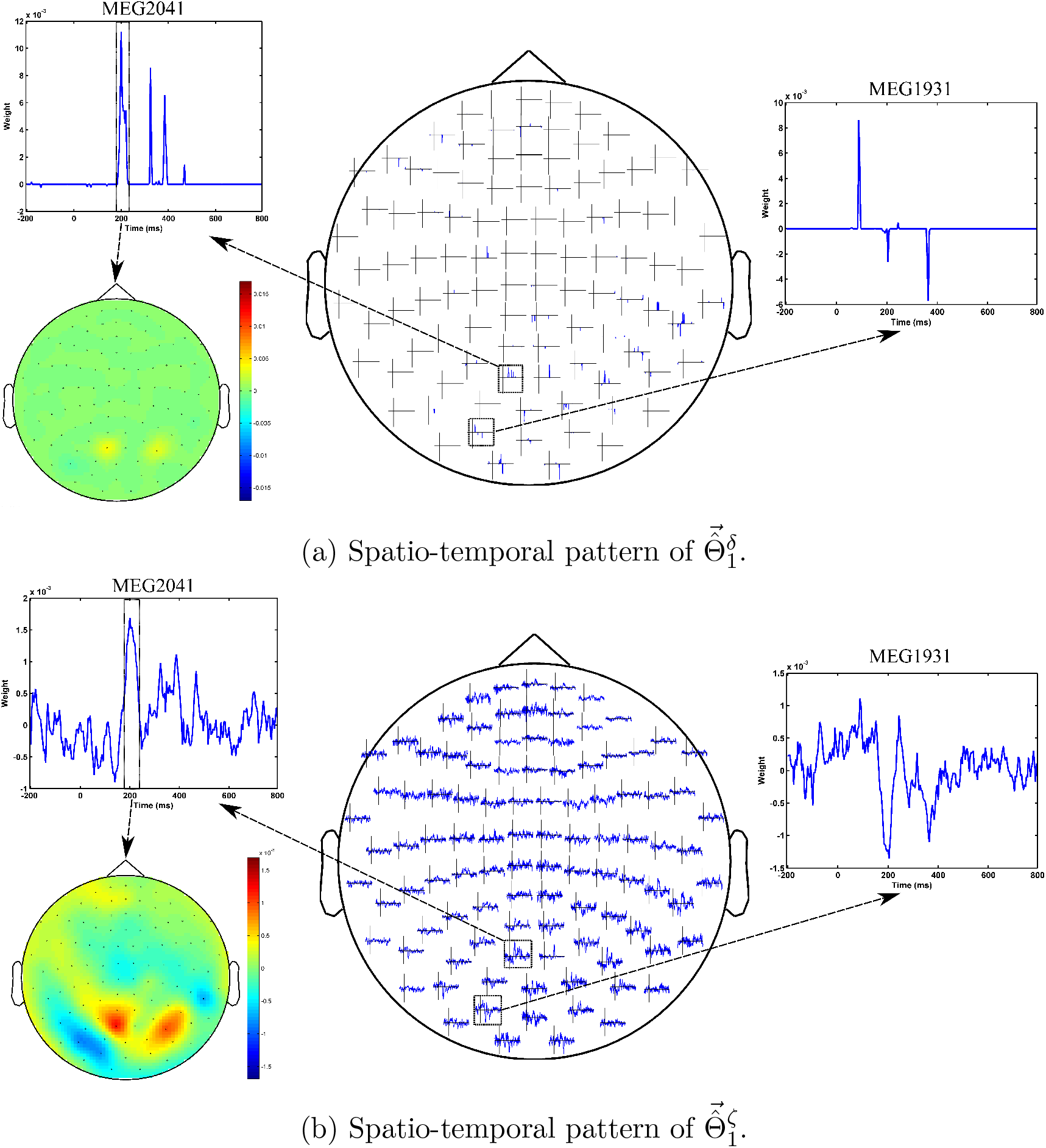
Comparison between spatio-temporal multivariate maps of the most accurate (5a) and the most interpretable (5b) classifiers for Subject 1. 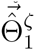 provides more spatio-temporal representativeness of the N170 effect than 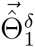.

**Figure 6.**
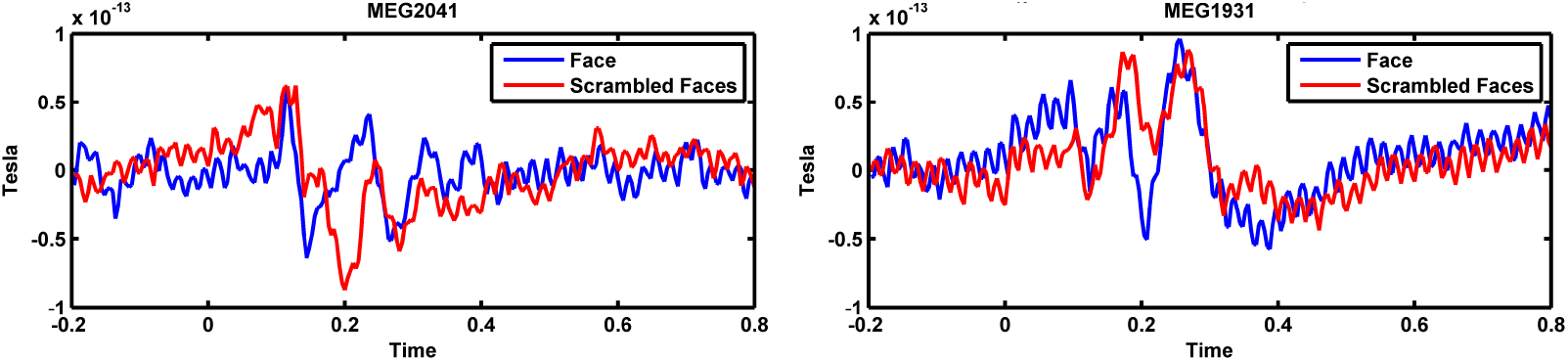
Event related fields (ERFs) of face and scrambled face samples for MEG2041 and MEG1931 sensors.

In addition, optimizing the hyper-parameters of brain decoding based on 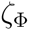 offers more reproducible brain decoders. According to table 2, using 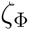 instead of 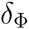 provides (on average) 0.15 more reproducibility over 16 subjects. To illustrate the advantage of higher reproducibility on the interpretability of maps, Figure 7 visualizes 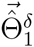 and 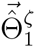 over 4 perturbed training sets. The spatial maps [Figure 7(a) and Figure 7(c)] are plotted for the magnetometer sensors averaged in the time interval between 184*ms* and 236*ms* after stimulus onset. The temporal maps [Figure 7(b) and Figure 7(d)] are showing the multivariate temporal maps of MEG1931 and MEG2041 sensors. While 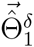 is unstable in time and space across the 4 perturbed training sets, 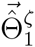 provides more reproducible maps.

**Figure 7.**
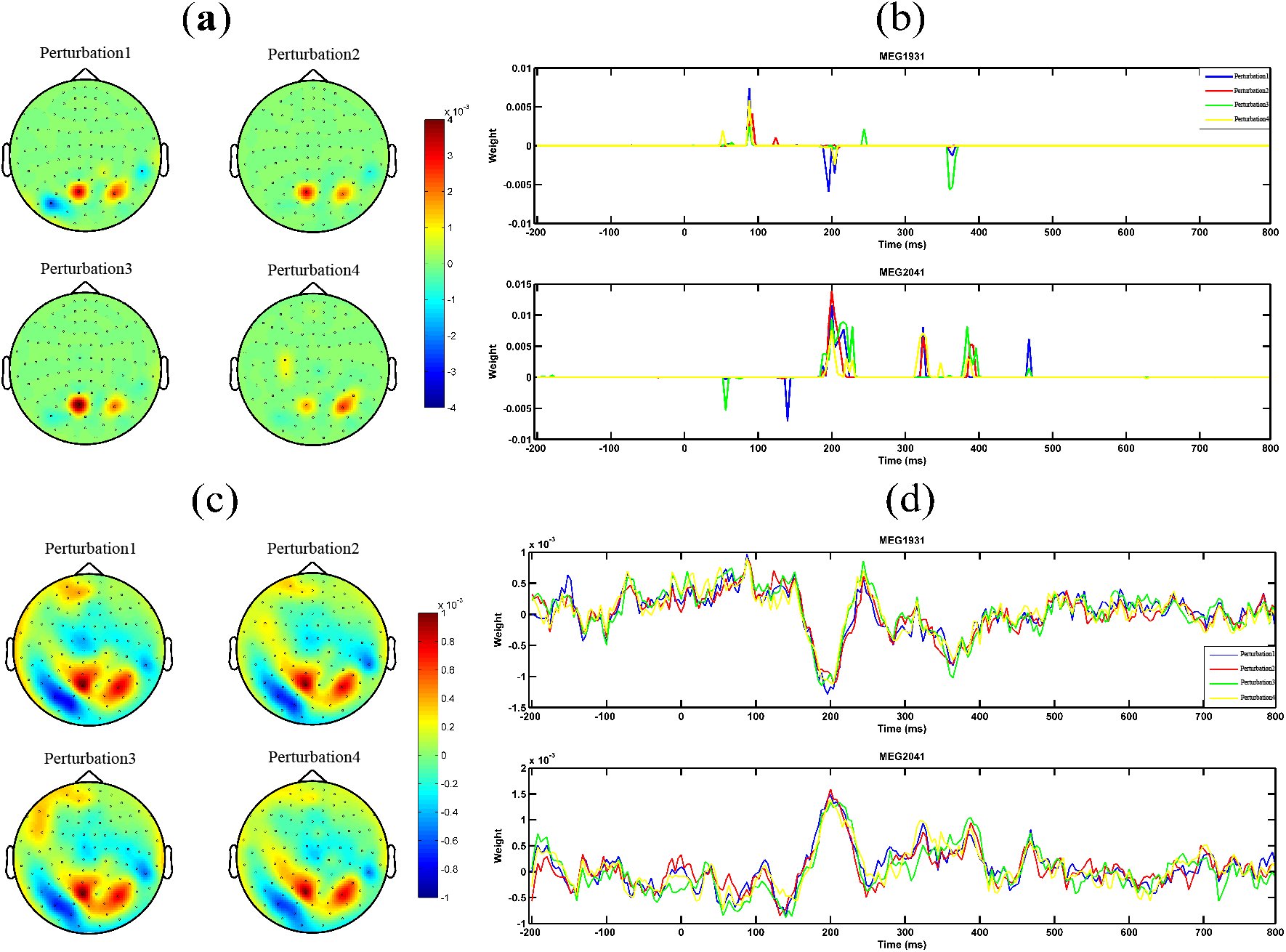
Comparison of the reproducibility of Lasso when 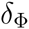 and 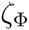 are used in the model selection procedure. (a) and (b) show the spatio-temporal patterns represented by 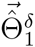 across the 4 perturbed training sets. (c) and (d) show the spatio-temporal patterns represented by 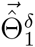 across the 4 perturbed training sets. Employing 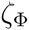 instead of 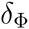 in the model selection yields more reproducible MBMs.

## 4 Discussions

### 4.1 Defining Interpretability: Theoretical Advantages

An overview of the brain decoding literature shows frequent co-occurrence of the terms interpretation, interpretable, and interpretability with the terms model, classification, parameter, decoding, method, feature, and pattern (see the quick meta-analysis on the literature in the supplementary material); however, a formal formulation of the interpretability is never presented. In this study, our primary interest is to present a theoretical definition of the interpretability of linear brain decoding models and their corresponding MBMs. Furthermore, we show the way in which interpretability is related to the reproducibility and neurophysiological representativeness of MBMs. Our definition and quantification of interpretability remains theoretical, as we assume that the true solution of the brain decoding problem is available. Despite this limitation, we argue that the presented definition provides a concrete framework of a previously abstract concept and that it establishes a theoretical background to explain an ambiguous phenomenon in the brain decoding context. We support our argument using an example in time-domain MEG decoding in which we show how the presented definition can be exploited to heuristically approximate the interpretability. This example shows how partial prior knowledge^1^ regarding underlying brain activity can be used to find more plausible multivariate patterns in data. Furthermore, the proposed decomposition of the interpretability of MBMs into their reproducibility and representativeness explains the relationship between the influential cooperative factors in the interpretability of brain decoding models and highlights the possibility of indirect and partial evaluation of interpretability by measuring these effective factors.

### 4.2 Application in Model Evaluation

Discriminative models in the framework of brain decoding provide higher sensitivity and specificity than univariate analysis in hypothesis testing of neuroimaging data. Although multivariate hypothesis testing is performed based solely on the generalization performance of classifiers, the emergent need for extracting reliable complementary information regarding the underlying neuronal activity motivated a considerable amount of research on improving and assessing the interpretability of classifiers and their associated MBMs. Despite ubiquitous use, the generalization performance of classifiers is not a reliable criterion for assessing the interpretability of brain decoding models [53]. Therefore, considering extra criteria might be required. However, because of the lack of a formal definition for interpretability, different characteristics of brain decoding models are considered as the main objective in improving their interpretability. Reproducibility [53, 54], stability selection [7, 47, 69], sparsity [96], and neurophysiological plausibility [97] are examples of related criteria.

Our definition of interpretability helped us to fill this gap by introducing a new multi-objective model selection criterion as a weighted compromise between interpretability and generalization performance of linear models. Our experimental results on single-subject decoding showed that adopting the new criterion for optimizing the hyper-parameters of brain decoding models is an important step toward reliable visualization of learned models from neuroimaging data. It is not the first time in the neuroimaging context that a new metric is proposed in combination with generalization performance for the model selection. Several recent studies proposed the combination of the reproducibility of the maps [53, 54, 43] or the stability of the classifiers [56, 57] with the performance of discriminative models to enhance the interpretability of decoding models. Our definition of interpretability supports the claim that the reproducibility is not the only effective factor in interpretability. Therefore, our contribution can be considered a complementary effort with respect to the state of the art of improving the interpretability of brain decoding at the model selection level.

Furthermore, this work presents an effective approach for evaluating the quality of different regularization strategies for improving the interpretability of MBMs. As briefly reviewed in Section 1, there is a trend in research within the brain decoding context in which prior knowledge is injected into the penalization term as a technique to improve the interpretability of decoding models. Thus far, in the literature, there is no ad-hoc method to compare these different methods. Our findings provide a further step toward direct evaluation of interpretability of the currently proposed penalization strategies. Such an evaluation can highlight the advantages and disadvantages of applying different strategies on different data types and facilitates the choice of appropriate methods for a certain application.

### 4.3 Regularization and Interpretability

Haufe et al. [39] demonstrated that the weight in linear discriminative models are unable to accurately assess the relationship between independent variables, primarily because of the contribution of noise in the decoding process. The problem is primarily caused by the decoding process that minimizes the classification error only considering the uncertainty in the output space [80, 98, 99] and not the uncertainty in the input space (or noise). The authors concluded that the interpretability of brain decoding cannot be improved using regularization. Our experimental results on the toy data (see Section 3.1) shows that if the right criterion is used for selecting the best values for hyper-parameters, appropriate choice of the regularization strategy can still play significant role in improving the interpretability of results. For example, in this case, the true generative function behind the sampled data is sparse (see Section 2.6.1), but because of the noise in the data, the sparse model is not the most accurate one. Using a more comprehensive criterion (in this case, 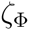) shows the advantage of selecting correct prior assumptions about the distribution of the data via regularization. This observation encourages the modification of the conclusion in [39] as follows: if the performance of the model is the only criterion in the model selection, then the interpretability cannot necessarily be improved by means of regularization.

### 4.4 Advantage over Mass-Univariate Analysis

Mass-univariate hypothesis testing methods are among the most popular tools in neuroscience research because they provide significance checks and a fair level of interpretability via univariate brain maps. Mass-univariate analyses consist of univariate statistical tests on single independent variables followed by multiple comparison correction. Generally, multiple comparison correction reduces the sensitivity of mass-univariate approaches because of the large number of univariate tests involved. Cluster-based permutation testing [5] provides a more sensitive univariate analysis framework by making the cluster assumption in the multiple comparison correction. Unfortunately, this method is not able to detect narrow spatio-temporal effects in the data [2]. As a remedy, brain decoding provides a very sensitive tool for hypothesis testing; it has the ability to detect multivariate patterns, but suffers from a low level of interpretability. Our study proposes a possible solution for the interpretability problem of classifiers, and therefore, it facilitates the application of brain decoding in the analysis of neuroimaging data. Our experimental results for the MEG data demonstrate that, although the non-parametric cluster-based permutation test is unable to detect the N170 effect in MEG data, employing 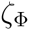 instead of 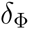 in model selection not only detects the stimuli-relevant information in the data, but also assures both reproducible and representative spatio-temporal mapping of the timing and the location of underlying neurophysiological effect.

### 4.5 Limitations and Future Directions

Despite theoretical and practical advantages, the proposed definition and quantification of interpretability suffer from some limitations. All of the presented concepts are defined for linear models, with the main assumption that 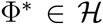 (where 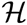 is a class of linear functions). This fact highlights the importance of linearizing the experimental protocol in the data collection phase [27]. Extending the definition of interpretability to non-linear models demands future research into the visualization of non-linear models in the form of brain maps. Currently, our findings cannot be directly applied to non-linear models. Furthermore, the proposed heuristic for the time-domain MEG data applies only to binary classification. One possible solution in mul-ticlass classification is to separate the decoding problem into several binary sub-problems. In addition the quality of the proposed heuristic is limited for the small sample size datasets (see supplementary material). Finding physiologically relevant heuristics for other acquisition modalities such as fMRI can be also considered in future work.

## 5 Conclusions

We presented a novel theoretical definition for the interpretability of linear brain decoding and associated multivariate brain maps. We demonstrated how the interpretability relates to the representativeness and reproducibility of brain decoding. Although it is theoretical, the presented definition provides a first step toward practical solution for filling the knowledge extraction gap in linear brain decoding. As an example of this major breakthrough, and to provide a proof of concept, a heuristic approach based on the contrast event-related field is proposed for practical evaluation of the interpretability in time-domain MEG decoding. We experimentally showed that adding the interpretability of brain decoding models as a criterion in the model selection procedure yields significantly higher interpretable models by sacrificing a negligible amount of performance. Our methodological and experimental achievements can be considered a complementary theoretical and practical effort that contributes to researches on enhancing the interpretability of mul-tivariate pattern analysis.

## Acknowledgments

The author wishes to thank Sandro Vega-Pons and Nathan Weisz for valuable discussions and comments.

## Appendix A. cERF and its Generative Nature

According to [39], for a linear discriminative model with parameters Θ, the unique equivalent generative model can be computed as follows:

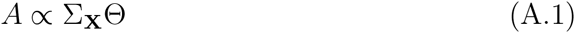

In a binary (**Y** = {1, -1}) least squares classification scenario, we have:

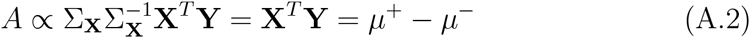

where Σ_X_ represents the covariance of the input matrix X, and *µ*^+^ and *µ*^-^are the means of positive and negative samples, respectively. Therefore, the equivalent generative model for the above classification problem can be derived by computing the difference between the mean of samples in two classes that is equivalent to the definition of cERF in time-domain MEG data.

**Figure B.8.**
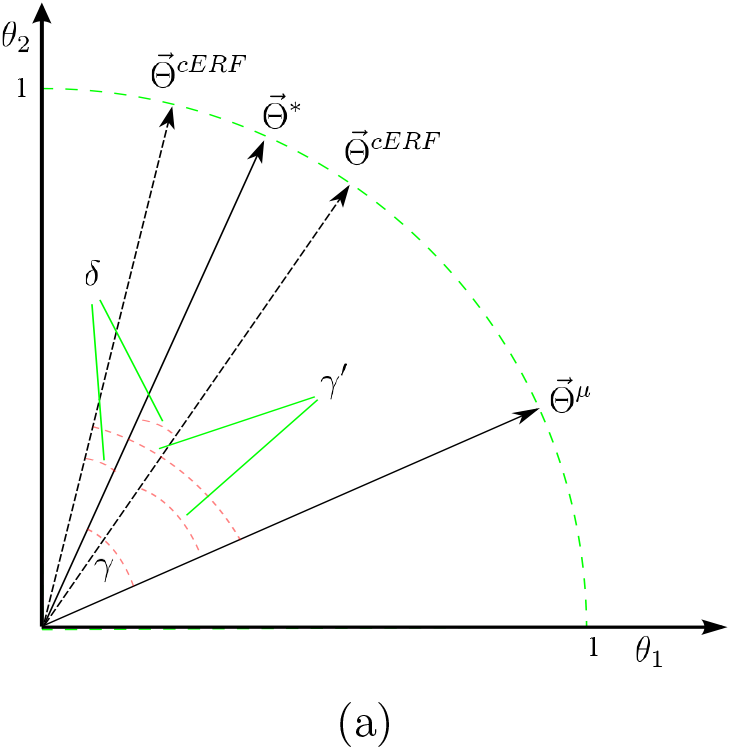
Misrepresentation of 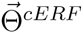 with respect to 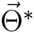.

## Appendix B. Relation between 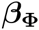 and 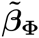 (Eq. 10)

Let γ be the angle between 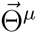 and 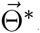. Let 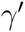 be the angle between 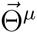 and 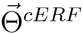. Furthermore, assume that δ is the angle between 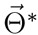 and 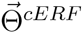 and that 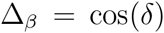. We consider both cases in which 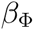 is underestimated/overestimated by 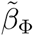 (see Figure B.8 as an example in 2-dimensional space). Then, we have:

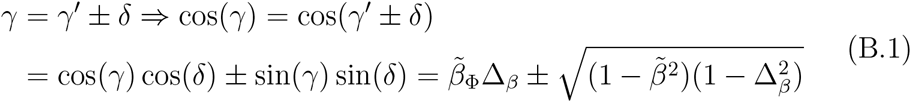

## Appendix C. Relation between 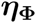 and 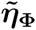 (Eq. 12)

Let 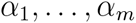 be the angles between 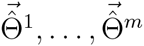 and 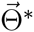, and 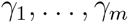 be the angles between 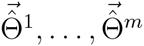 and 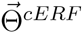. Furthermore, assume that

**Figure C.9.**
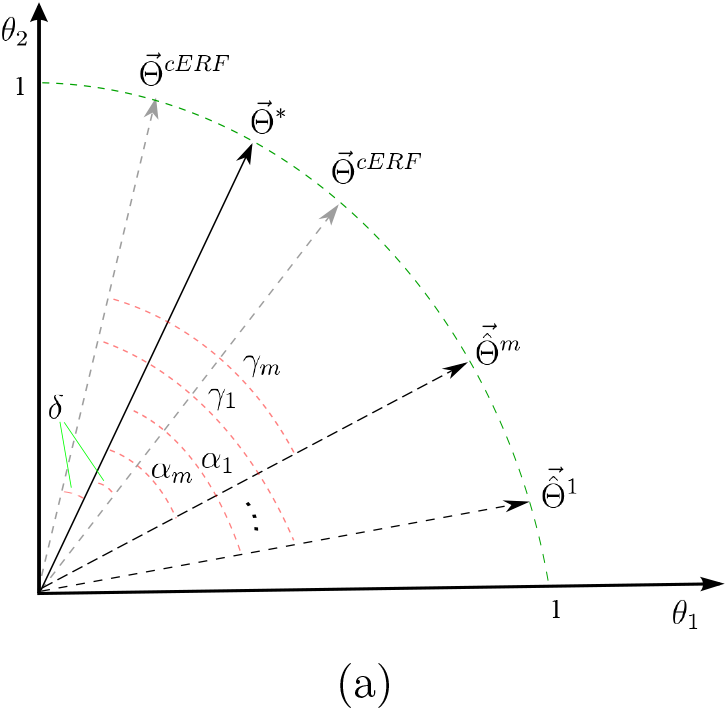
Relation between 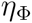 and 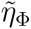.

*δ* is the angle between 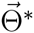 and 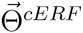. We consider both cases in which 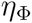 is underestimated/overestimated by 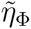 (see Figure C.9 as an example in 2-dimensional space).

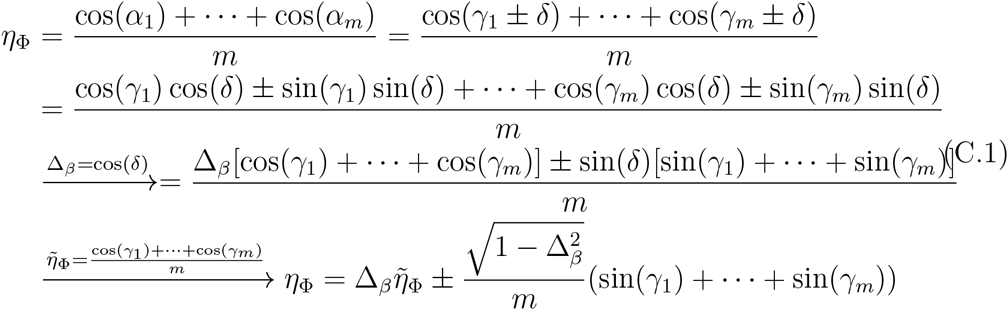

## Appendix D. Proof of Proposition 1

Throughout this proof, we assume that all of the parameter vectors are normalized in the unit hypersphere (see Figure D.10 as an illustrative example in 2 dimensions). Let 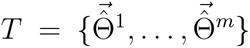 be a set *m* MBMs, for *m* perturbed training sets where 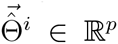. Now, consider any arbitrary *p -* 1-dimensional hyperplane 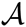 that contains 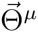. Clearly, 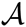 divides the p-dimensional parameter space into 2 subspaces. Let 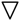 and 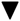 be binary

operators where 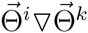 indicates that 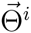 and 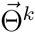 are in the same subspace, and 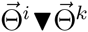 indicates that they are in different subspaces. Now, we define 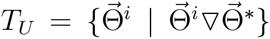 and 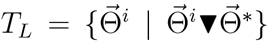. Let the cardinality of *T_L_* denoted by *n*(*T*_L_) be *j* (*n*(*T_L_*) = *j*). Thus, *n*(*T_U_*) = *m - j*. Now, assume that 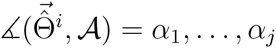 are the angles between 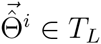 and 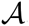, and (similarly) 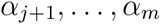 for 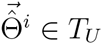 and 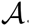. Based on Eq. 5, let 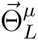 and 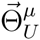 be the main maps of *T_L_* and *T_U_*, respectively. Therefore, we obtain 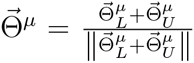 and 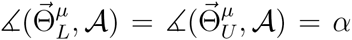. Furthermore, assume 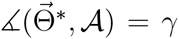. As a result, 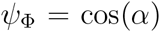 and 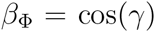. According to Eq. 4 and using a cosine similarity definition, we have:

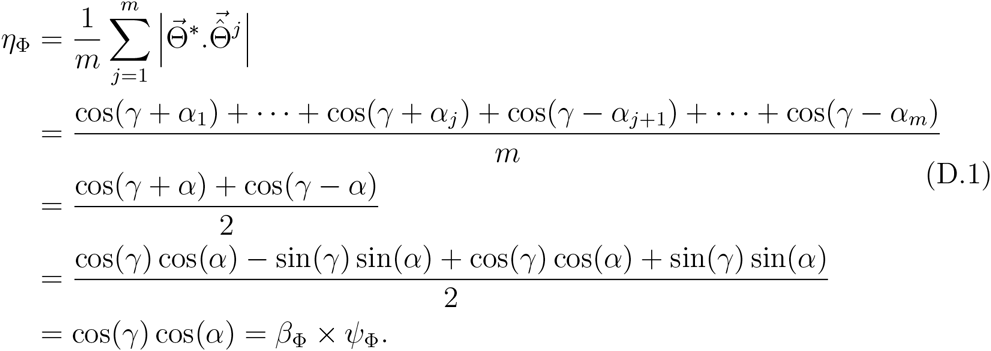

A similar procedure can be used to prove 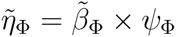 by replacing 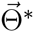 with 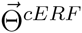.

## Appendix E. Computing the Bias and Variance in Binary Classification

Here, using the out-of-bag (OOB) technique, and based on procedures proposed by [83] and [100], we compute the expected prediction error (EPE) for a linear binary classifier Φ under bootstrap perturbation of the training set. Let *m* be the number of perturbed training sets resulting from partitioning (*X*,*Y*) into (*X_tr_*,*Y_tr_*) and (*X_ts_*,*Y_ts_*), i.e., training and test sets. If 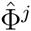 is the linear classifier estimated from the jth perturbed training set, then the main prediction 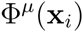 for each sample in the dataset can be computed as follows:

**Figure D.10.**
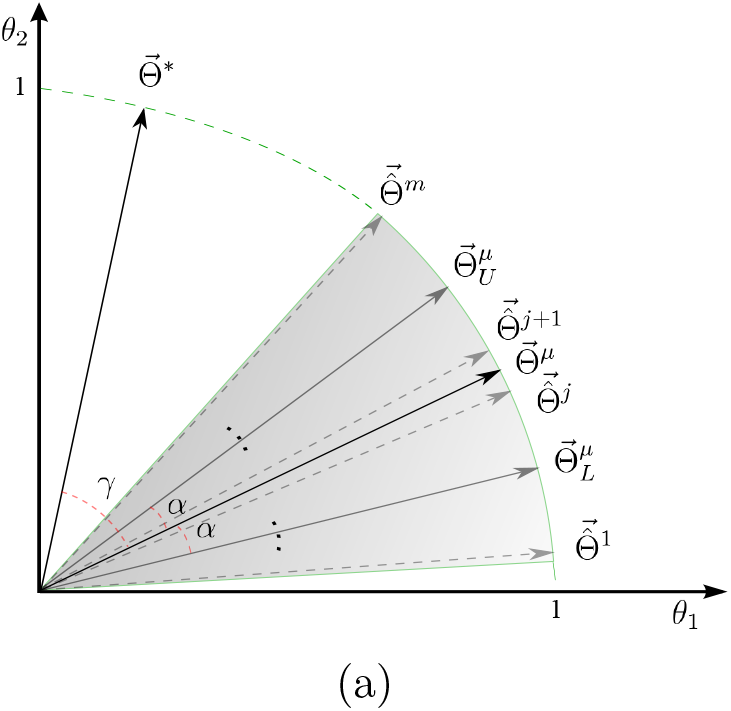
Relation between representativeness, reproducibility, and interpretability in 2 dimensions.

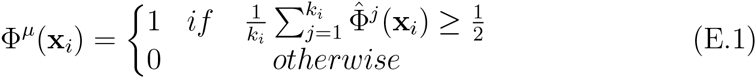

where *k*_i_ is the number of times that *x_i_* is present in the test set^1^.

The computation of bias is challenging because the optimal model 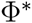 is unknown. According to [101], misclassification error is one of the loss measures that satisfies a Pythagorean-type equality, and:

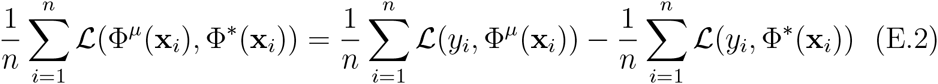

Because all terms of the above equation are positive, the mean loss between the main prediction and the actual labels can be considered as an upper-bound for the bias:

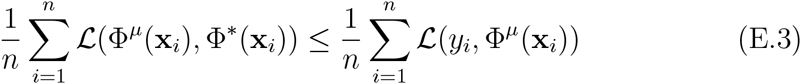

Therefore, a pessimistic approximation of bias *B*(*X_i_*) can be calculated as follows:

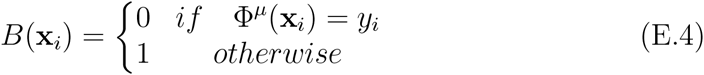

Then, the unbiased and biased variances (see [83] for definitions) in each training set can be calculated by:

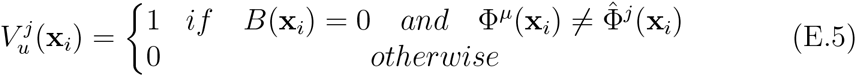

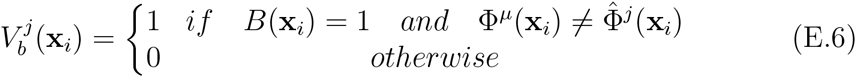

Then, the expected prediction error of Φ can be computed as follows (ignoring the irreducible error):

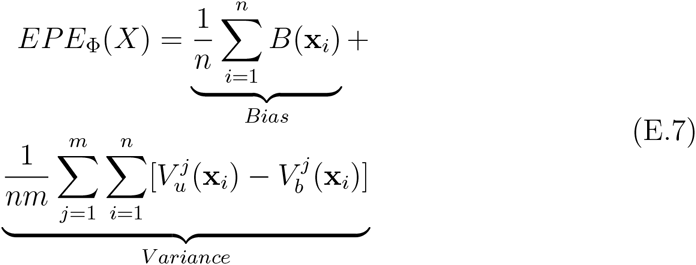

## Footnotes

^1^Here, neurophysiological plausibility refers to the spatio-temporal chemo-physical constraints of the underlying neural activity that is highly dependent on the acquisition device.

^1^The application of the presented heuristic to MEG data can be extended to EEG because of the inherent similarity of the measured neural correlates in these two devices. In the EEG context, the ERF can be replaced by the event-related potential (ERP).

^1^The full dataset is publicly available at ftp://ftp.mrc-cbu.cam.ac.uk/personal/rik.henson/wakemandg_hensonrn/

^2^The competition data are available at http://www.kaggle.com/c/decoding-the-human-brain

^1^The preprocessing scripts in python and MATLAB are available at: https://github.com/FBK-NILab/DecMeg2014/

^2^The MATLAB code used for experiments is available at https://github.com/smkia/interpretability/

^1^The bounds of colorbars are symmetrized based on the maximum absolute value of parameters

^1^The partial knowledge can be based on already known facts regarding the timing and location of neural activity.

^1^It is expected that each sample x_i_ ε *X* appears (on average) 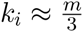 times in the test sets.

